# Identification of stress specific autophagy regulators from tandem CRISPR screens

**DOI:** 10.1101/2024.03.27.587008

**Authors:** Truc T. Losier, Maxime W.C. Rousseaux, Ryan C. Russell

## Abstract

Autophagy is a conserved degradative process that promotes cellular homeostasis under stress conditions. Under nutrient starvation autophagy is largely non-selective, promoting the indiscriminate breakdown of cytosolic components. Conversely, selective autophagy is responsible for the specific turnover of damaged organelles including endoplasmic reticula, lysosomes, mitochondria, and peroxisomes. The mechanisms of selective autophagy are best understood through the activity of cargo-specific receptors called autophagy receptors, which facilitate the engulfment of the targeted cargo within autophagosomes, leading to subsequent degradation. We hypothesized that selective autophagy may be regulated by distinct upstream signaling from starvation induced autophagy, providing an additional layer of regulatory control to targeted autophagic degradation. To comprehensively address this question we conducted kinome-wide CRISPR screens to identify distinct signaling pathways responsible for the regulation of basal autophagy, starvation-induced autophagy, and two types of selective autophagy, ER-phagy and pexophagy. These parallel screens identified established and novel autophagy shared regulators under these conditions, as well as kinases specifically required for ER-phagy or pexophagy. More specifically, CDK11A and NME3 were further characterized to be selective ER-phagy regulators. Meanwhile, PAN3 and CDC42BPG were identified as activator or inhibitor of pexophagy, respectively. Collectively, these datasets provide the first comparative description of the kinase signaling specificity, separating regulation of selective autophagy and bulk autophagy.

**Highlights:**

- Parallel pooled kinome genetic knockout screens reveal known and novel regulators of autophagy under basal conditions, nutrient starvation, ER stress, and peroxisomal stress
- Selective ER and peroxisomal autophagy both have unique activators and inhibitors that distinguish them from bulk autophagy
- CDK11A and NME3 specifically induce and inhibit ER-phagy, respectively
- PAN3 and CDC42BPG specifically induce and inhibit pexophagy, respectively

## Introduction

Macroautophagy (hereafter referred to as autophagy) is a cellular degradative program that is upregulated by cellular stressors^1^. The autophagy pathway is driven by the formation of a double membrane vesicle called an autophagosome that sequesters cytosolic cargo for degradation^1^. Degradation of autophagy cargo is achieved through the activity of hydrolases that are supplied by lysosomes after their fusion to autophagosomes^2^. Autophagosome formation is mediated through the activity of a conserved group of autophagy-related (ATG) proteins^1^. Autophagosome biogenesis is promoted by the activity of a serine/threonine kinase call ULK1^3^. ULK1 phosphorylates and activates multiple ATG proteins, including components of the lipid kinase VPS34 to promote stress-induced autophagy^4,5,6,7,8,9^. Maturation of the autophagosome and sequestration of targeted cargo require the lipidation of ATG8 family members (LC3A, B and C, GABARAP, GABARAPL1 and GABARAPL2) to the lipid phosphatidylethanolamine^10^. LC3B is the best studied member of the ATG8 family in mammals^11,12^. Autophagy was originally described as a bulk degradation pathway that indiscriminately engulfs and degrades cytosolic components^13^. However, the importance of autophagy as a targeted degradation pathway has been established in normal and disease biology^14,15^.

The targeted degradation of cargo by the autophagy pathway is called selective autophagy^16,17^. Selective autophagy can be categorized into various subgroups based on the specific cellular components targeted for degradation: degradation of mitochondria (mitophagy), peroxisomes (pexophagy), endoplasmic reticulum (ER-phagy), lipid (lipophagy), lysosomes (lysophagy), ribosomes (ribophagy), aggregated proteins (aggrephagy), and pathogens (xenophagy)^17,18,19,20,21,22,23^. Selective autophagy requires all the core ATG proteins of bulk autophagy, but also includes a class of proteins called autophagy receptors^24,25^. Autophagy receptors bind damaged or harmful intracellular components and recruit them to autophagosomes for degradation through LC3 interaction^17^. Autophagy receptors can be divided into ubiquitin-bound and membrane-associated classes^17,26^. Well-known ubiquitin-bound receptors are SQSTM1/p62, NBR1, NDP52, and OPTN^17^. They typically contain both LC3-interacting region (LIR) and ubiquitin-binding domains (UBD), which allow them to recruit ubiquitinated targets to autophagosomes for degradation^17^. Membrane-bound receptors localize, or are constitutively present on the target organelles and are responsible for damaged organelle recognition by autophagy machinery^27^. Importantly, defective selective autophagy has been implicated in several human diseases^28,29^. For instance, impaired clearance of ER is linked to neurodegenerative disorders such as Alzheimer’s disease and amyotrophic lateral sclerosis due to the accumulation of misfolded proteins and ER stress, contributing to neuronal damage^30^. Defective pexophagy is linked to the pathology of Zellweger spectrum disorders^31,32^. Additionally, impaired pexophagy results in the accumulation of dysfunctional peroxisomes and subsequently exacerbation of liver damage, which contributes to liver diseases such as non-alcoholic fatty liver disease and alcoholic liver disease^33^. Dysregulation in lysophagy contributes to infection and cancer^34^. Impaired xenophagy is linked to inflammatory bowel diseases and chronic infectious diseases including tuberculosis^35,36,37^.

In this study, we chose ER-phagy and pexophagy as our models of selective autophagy since they have not been characterized as extensively as other forms of selective autophagy like mitophagy or xenophagy, making them ideal candidates for comprehensive analysis. Selective autophagic degradation of the ER is a homeostatic mechanism, maintaining ER size, removal of aggregated or miss-folded proteins, and turnover of ER damage^38^. To date, six ER-phagy receptors have been identified in mammals, including FAM134B, RTN3L, CCPG1, SEC62, TEX264, and ATL3^39,40,41,42,43,44,45^. These receptors localize to ER sub-compartments and are capable of recruiting autophagy machinery to the ER through LC3/GABARAP-interacting regions (LIR/GIM)^46^. For instance, FAM134B primarily facilitates the degradation of sheet ER, while ATL3 and RTN3L mediate degradation of tubular ER^39,41,45^. FAM134B and RTN3L have been reported to induce ER fragmentation through reticulon homology domains (RHD), which contributes to ER turnover^41,45^. SEC62 plays an important role in recovery from ER stress by promoting autophagic clearance of select ER regions through LIR domain^44^. Some of the receptors have been shown to regulate ER-phagy through interaction with upstream autophagy complexes in addition to autophagosomal LC3/GABARAP family proteins. For example, CCPG1 can bind directly to FIP200, a component of ULK1 complex^40^. TEX264 has also been reported to interact with proteins associated with an early phase of autophagosome formation, including FIP200 and WIPI2^47^. In addition to autophagy receptors, new pathways have been reported to positively regulate ER-phagy^48^. Perturbation of mitochondrial oxidative phosphorylation system hinders ER-phagy^48^. DDRGK1 has been shown to recruit UFMylation (ubiquitin-like post-translational modification) machinery to the ER surface, which is critical for autophagic degradation of ER sheets^48^.

Peroxisomes are small single membrane-bound organelles important for synthesis and metabolism of reactive oxygen species, and the alpha- and beta-oxidation of branched chain and very long chain fatty acids (VLCFA)^49,50,51^. Peroxisome homeostasis is tightly regulated and is achieved by a set of peroxin (PEX) proteins^19^. Stress conditions such as starvation, hypoxia, or incubation with reactive oxygen species (ROS) disturb peroxisome homeostasis and result in autophagic degradation of peroxisomes^52,53,54^. Peroxisomes require proper import of matrix proteins to become functional^19^. Therefore, the significance of the matrix import system for peroxisome function may underscore its role in peroxisome quality control. Indeed, several components of the matrix import systems have been linked to pexophagy regulation, such as PEX2, PEX5, PEX13, and PEX14^19,55^. Under stress, the peroxisomal E3 ubiquitin ligase PEX2 is activated and ubiquitinates peroxisomal proteins PMP70 and PEX5^53,54^. Ubiquitinated peroxisomes are then recruited to autophagosomal sites for degradation through interactions with the autophagy receptors NBR1 and p62^53,54^. PEX14 has been shown to preferentially bind to LC3 to promote pexophagy during starvation^56^. PEX13 levels are reduced during the early timepoints of starvation to promote pexophagy induction^55^. Additionally, elevated ROS levels and hypoxia trigger different cellular responses, which subsequently enhances PEX5 ubiquitination and induces pexophagy^19,54,57^. Conversely, ubiquitinated peroxisome proteins are removed from peroxisomal membrane by the AAA-type ATPase PEX1-PEX6-PEX26 and the deubiquitinase USP30, which hinders pexophagy^58,59,60^.

Kinase-mediated phosphorylation plays in important role in autophagy initiation. Induction of the autophagy pathway is best characterized in the context of nutrient starvation, where nutrient-sensitive kinases, including mTORC1 and AMPK, are critical for modulating autophagy activation^61,62,63^. Under nutrient sufficiency mTORC1 suppresses autophagy through direct phosphorylation of several components associated with autophagy induction^64^. mTORC1-mediated autophagy regulation is also achieved through direct inhibitory phosphorylation of ULK1^65,66,67^. Upon starvation, mTORC1 is inactivated, which results in ULK1 release from inhibitory phosphorylation and subsequent autophagy induction^68^. AMPK activity under energy and nutrient starvation can regulate autophagy through modulation of ULK1 and mTORC1 activity^65,68,69^. In addition, AMPK and mTORC1 have been reported to directly regulate the activity of the VPS34 kinases, ensuring a precisely controlled initiation of autophagy in response to various cellular stresses^70,71^. Collectively, these nutrient-dependent kinases play an essential role in mediating the activation of bulk autophagy in response to starvation.

Several genome- and kinome-wide screens have been performed to identify basal autophagy regulators using RNAi or CRISPR/Cas9-based targeting^72,73,74,75,76,77^. Additionally, starvation-induced autophagy screens have been conducted to establish autophagy regulators under nutrient stress pathways^3,78,79^. Focus has more recently shifted towards selective autophagy regulation. For example, a genome wide screen using an ER-phagy specific reporter identified the UFMylation pathway as a mediator of ER-phagy, while a screen for regulators of PARKIN stability identified genes implicated in mitophagy^48,80^. A limitation in the approaches employed above is the difficulty that arises when attempting to compare these independent datasets, which each tend to look at one facet of autophagy. This is particularly important when comparing the signaling that might underly selective vs. bulk autophagy. We envisioned that novel kinases may play a key role in cargo specific autophagy, like how mTORC1 and AMPK are critical to starvation-induced autophagy. To address this gap, we used an autophagy receptor, p62, which has been implicated in several types of selective autophagy as a common reporter. For tight experimental control and broad comparison across stressors, we performed pooled kinome wide CRISPR screens from the same polyclonal cell population varying only by the acute stress conditions. Using this approach, we identified activators and inhibitors of ER-phagy and pexophagy, which differentiated them from basal or starvation induced autophagy. The datasets described in this study serves as a benchmark resource that not only represents a comprehensive analysis of bulk autophagy, but also provides a unique look at the signaling specificity underlying ER-phagy and pexophagy.

## Results

### A kinome-scale CRISPR screen using an autophagic flux reporter

To measure autophagy rates in a high throughput, quantitative manner, we first generated a stable cell line with a fluorescent marker of autophagy flux, p62/SQSTM1. p62 is capable of binding to multiple types of ubiquitinated cargo and recruiting them to autophagosomes for degradation. p62 was chosen for this study because it is involved in nearly all forms of selective autophagy and can be used as a marker for autophagic flux for several types of selective autophagy. To measure changes in autophagic flux, we constructed a dual-fluorescence reporter, DsRed-IRES-GFP-p62 (Figure 1A). HEK293A cells were stably transduced with constructs expressing GFP-tagged p62 with an internal DsRed control (DsRed-IRES-GFP-p62, Figure 1A). From the polyclonal reporter line, we generated and assessed responses of monoclonal populations to amino acid-free media, a potent stress condition to induce autophagy. In this system, flow cytometry of fluorescence-activated cell sorting (FACS) allows for the rapid and quantitative measurement of relative p62 levels (GFP) while controlling for non-selective changes in protein abundance (DsRed control, Figure 1A, right panel). Levels of p62 decrease during autophagy activation, resulting in a left shift in the GFP fluorescence signal (Figure 1A, right panel). Conversely, autophagy inhibition leads to a right shift in the GFP signal (Figure 1A, right panel). We benchmarked our reporter line against known p62 responses using both western blot (WB) and flow cytometry, identifying ideal timepoints and concentrations for measuring stress-based p62 flux. The DsRed-IRES-GFP-p62 reporter line showed a robust clearance of tagged p62 in response to amino acid starvation, with no changes in DsRed control (Figure 1B, left panel). We selected a 3-hour period of starvation for screening purposes, as it was the earliest timepoint giving peak p62 degradation. While the induction of autophagy by starvation was clear by western blot analysis, we next sought to verify whether that stress condition triggered similar responses by FACS analysis. Consistently, we observed a significant reduction in GFP signal while there was no change in DsRed fluorescence in the starved reporter cells compared to the untreated ones (Figure 1B, right panel). We next investigated the autophagic response to ER stress and peroxisomal stress. ER stress was induced by tunicamycin treatment using established ranges and concentrations^81,82^. Tunicamycin hinders the first stage of N-linked glycan production in proteins and leads to the accumulation of improperly folded proteins. Compared to the unstimulated cells, we observed robust autophagic flux in tunicamycin-treated cells under all examined conditions, as shown through substantial reductions in p62 levels (Figure 1C). We selected 10 µg/mL of tunicamycin for a duration of 6 hours as the designated condition to induce ER-phagy for our screening experiments. This was further confirmed using flow cytometry, where exposure to tunicamycin produced a significant decrease in GFP-p62 fluorescence compared to the untreated cells, with no changes in DsRed (Figure 1C, right panel). We then characterized the optimal timepoints for inducing p62 flux following peroxisomal stress using both WB and flow cytometry (Figure 1D). To initiate peroxisomal stress, we treated cells with clofibrate to disrupt peroxisome function and induce pexophagy^83^. Using previously reported timepoints and concentrations, we observed a decrease in p62 levels in all clofibrate-treated samples with no changes in DsRed, indicating an activation of autophagy (Figure 1D). Among the conditions tested, the induction of peroxisomal stress using clofibrate (1 mM) for a duration of 6 hours elicited a robust and consistent response in the reporter cell line via both orthogonal approaches (Figure 1D). For each of the identified stressors, we also confirmed that the effects on p62 flux was due to autophagy by repeating the stress analysis in an autophagy-deficient background (ATG5 KO, DsRed-IRES-GFP-p62 293A cells, Figure S1). In all stress conditions, we found that the stress-induced decrease in GFP-p62 fluorescence was blocked in autophagy deficient cells, indicating stress induced p62 clearance optimized above was a result of autophagic flux. Collectively, these experiments have established the suitability of our reporter cell line, stress conditions, and screening methodology to discover novel kinases that regulate selective autophagy.

**Figure 1:**
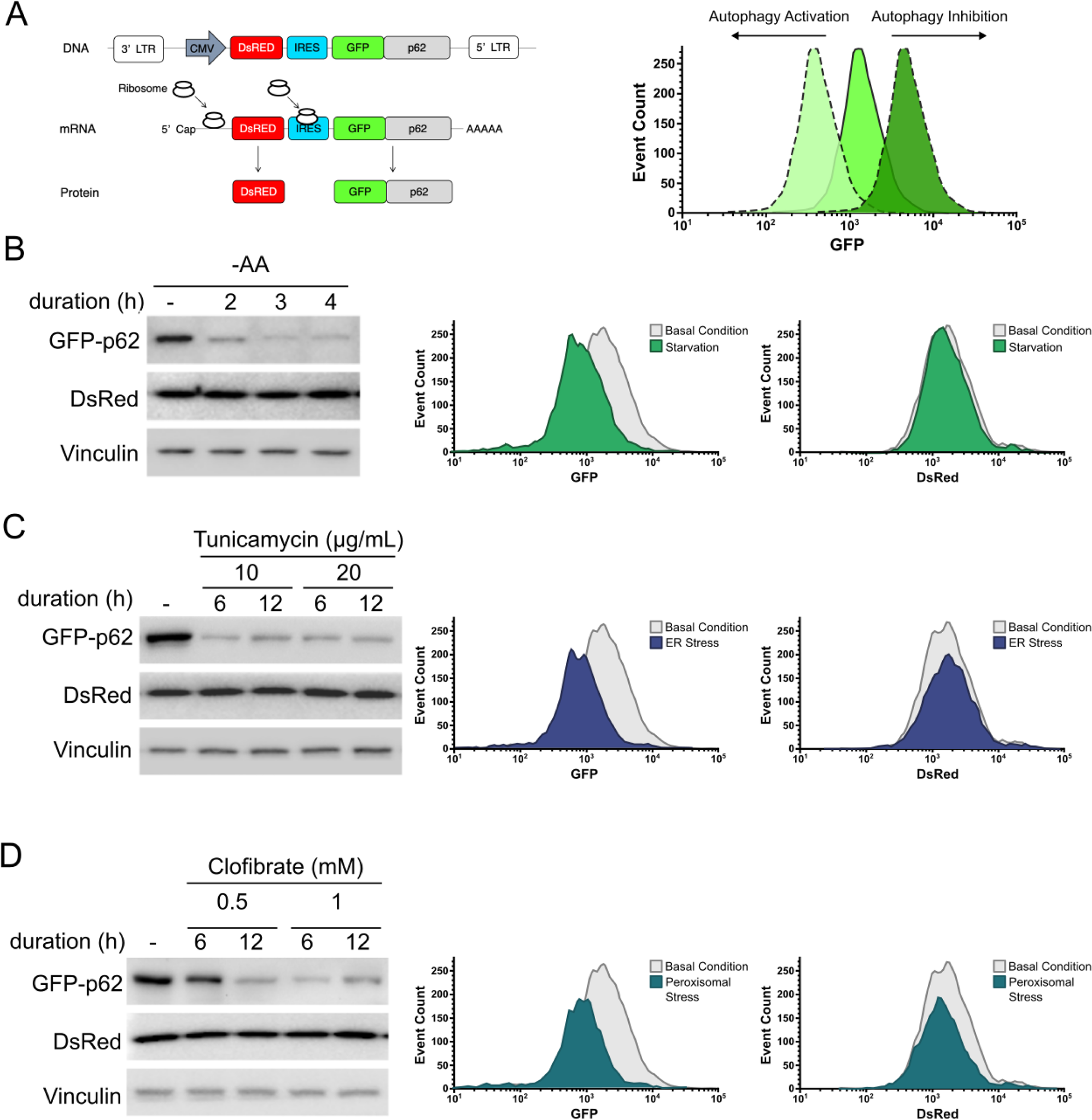
CRISPR-based kinome-wide screen using an autophagic flux reporter. (A) Schematic representation of the autophagic flux reporter DsRed and GFP-tagged p62. p62 is selectively incorporated into and degraded along with the autophagosomal membrane. Therefore, the expression level of GFP-p62 is inversely proportional to autophagic flux. Examples of autophagy activation and inhibition were demonstrated using the histogram in the right panel. (B) The reporter cell line was treated with starvation in time- and concentration-dependent manners. WB was used to examine DsRed and GFP-p62 signals. FACS was employed to investigate GFP and DsRed fluorescence of the reporter cells treated with amino acid-free media for 3 hours. Histograms were used to depict changes in GFP-p62 and DsRed levels. (C) The reporter cell line was incubated with tunicamycin in time- and concentration-dependent manners. DsRed and GFP-p62 were analysed using WB. FACS was used to examine GFP and DsRed signals of the reporter cells exposed to tunicamycin (10 µg/mL) for 6 hours. Histograms were used to depict changes in GFP-p62 and DsRed levels. (D) The reporter cells were incubated with clofibrate in time- and concentration-dependent manners. DsRed and p62 levels were examined using WB. FACS was employed to investigate GFP and DsRed fluorescence of the reporter cells treated with clofibrate (1 mM) for 6 hours. Histograms were used to depict changes in GFP-p62 and DsRed levels.

### CRISPR-based screens identify shared and distinct of stress-selective autophagic pathways

To gain insight into signaling pathways underlying selective autophagic flux, we conducted kinome-wide CRISPR-based screens using the validated DsRed-IRES-GFP-p62 reporter line (Figure 2A). Briefly, cells were transduced with the Brunello kinome library, a pooled lentiCRISPRv2 library containing 3,052 unique sgRNAs targeting 763 human kinases genes (4 guides per target) at a multiplicity of infection of 0.3 to minimize dual viral integration^84^. Infected cells were selected for 3 days with puromycin (1 µg/mL), and one quarter of the infected cells were subsequently exposed to either amino acid-free DMEM media (for 3 hours; starvation), tunicamycin (at 10 μg/mL, 6 hours; ER stress), clofibrate (at 1 mM, 6 hours; peroxisomal stress), or were left untreated (basal) on the 16th day post-viral transduction. The untreated, starvation-treated, tunicamycin-treated, and clofibrate-treated cells were fixed with 2 % PFA and harvested on the same day to allow for direct comparison among stress conditions from the population of edited cells. Fixed samples were analysed by FACS. Approximately 10 % gated populations were selected in the high (to identify activators) and low (to identify inhibitors) GFP/DsRed regions (Figure 2A). Genomic DNA from the fixed sorted cells, as well as the corresponding fixed pre-sort population, which controls for alternations in sgRNA abundance mediated by changes in cellular fitness, were extracted and sgRNA sequences were amplified and analysed by next generation sequencing (NGS). The screen was performed 4 times, each of which included all 4 conditions, for a total of 16 individual kinome screens.

**Figure 2:**
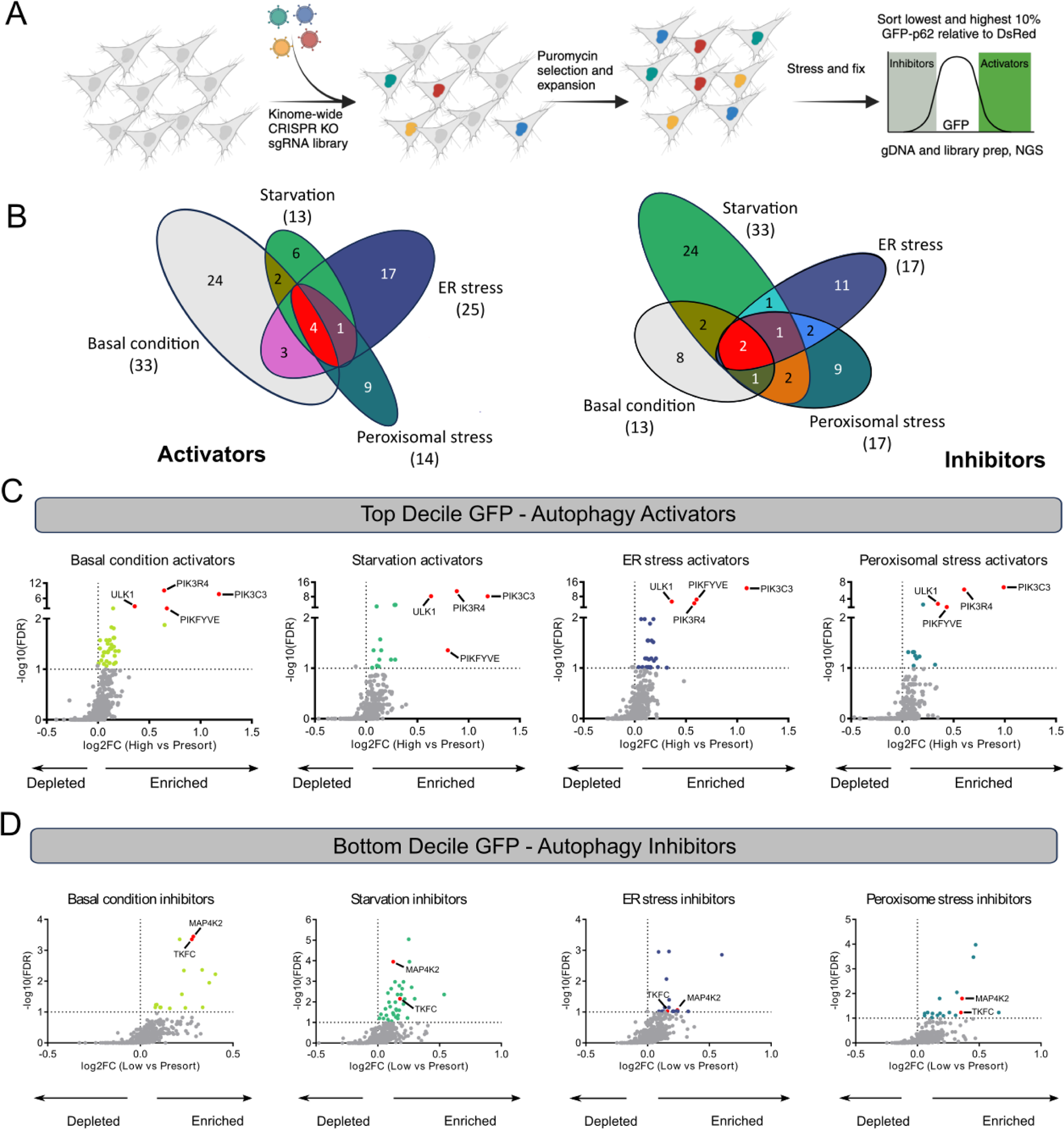
CRISPR-based screens identify known regulators in core autophagy machinery and several genes involved in selective autophagy pathways. (A) The screening strategy used to identify positive regulators and negative regulators of autophagy pathways. (B) Venn diagrams depict shared and distinct activators or inhibitors among 4 examined conditions. (C), (D) Volcano plots from the four screens. For each gene, the x-axis indicates its enrichment or depletion, determined by the mean of all four sgRNAs targeting the gene, in the sorted population compared to the corresponding unsorted population. The y-axis represents the statistical significance, as indicated by the false discovery rate (FDR)-corrected p-value. The horizontal dashed line represents an FDR-value threshold of 0.1. Red dots on the graph denote common regulators across all four conditions, hits specific to different conditions are colored accordingly, and all other genes are represented as gray dots.

We conducted analysis of sgRNA enrichment and depletion between the sorted populations and the corresponding unsorted populations across experimental replicates for each stress condition using CRISPRBetaBinomial^85^. Hits were called using false discovery rate (FDR; adjusted *p* values) and a positive log_2_ fold change (log_2_FC); detailed in Table 1. Log_2_FC represents enrichment or depletion of each gene, measured by the mean of all four sgRNAs targeting the gene, in the sorted high or low GFP/DsRed population compared to the corresponding unsorted population. FDR indicates the statistical significance of sgRNAs abundance of a gene in the sorted high or low GFP/DsRed group compared to the unsorted sample. Overlap between autophagy activators and inhibitors among 4 conditions showed some shared regulators, but also a large degree of unique hits (Figure 2B). Volcano plots showcasing FDR and log_2_FC values were constructed to visualize autophagy activators and inhibitors (Figures 2C and 2D). The asymmetric nature of the plot suggests that the statistical ability to detect sgRNA enrichment (positive log_2_FC) was greater than the ability to detect sgRNA depletion (negative log_2_FC) in the sorted population, a feature that has been previously reported^86^. Therefore, subsequent analyses focused exclusively on genes with FDR <0.1 for enrichment (positive log_2_FC) in the sorted population. Additionally, hits were omitted if they possessed significant FDR values in both positive and negative regulator populations (conflicts)^87^. A summary of common and distinct regulators that meet the criteria of FDR cutoff and positive log_2_FC from both autophagy-activating and inhibiting populations are provided in Table 1.

**Table 1:**
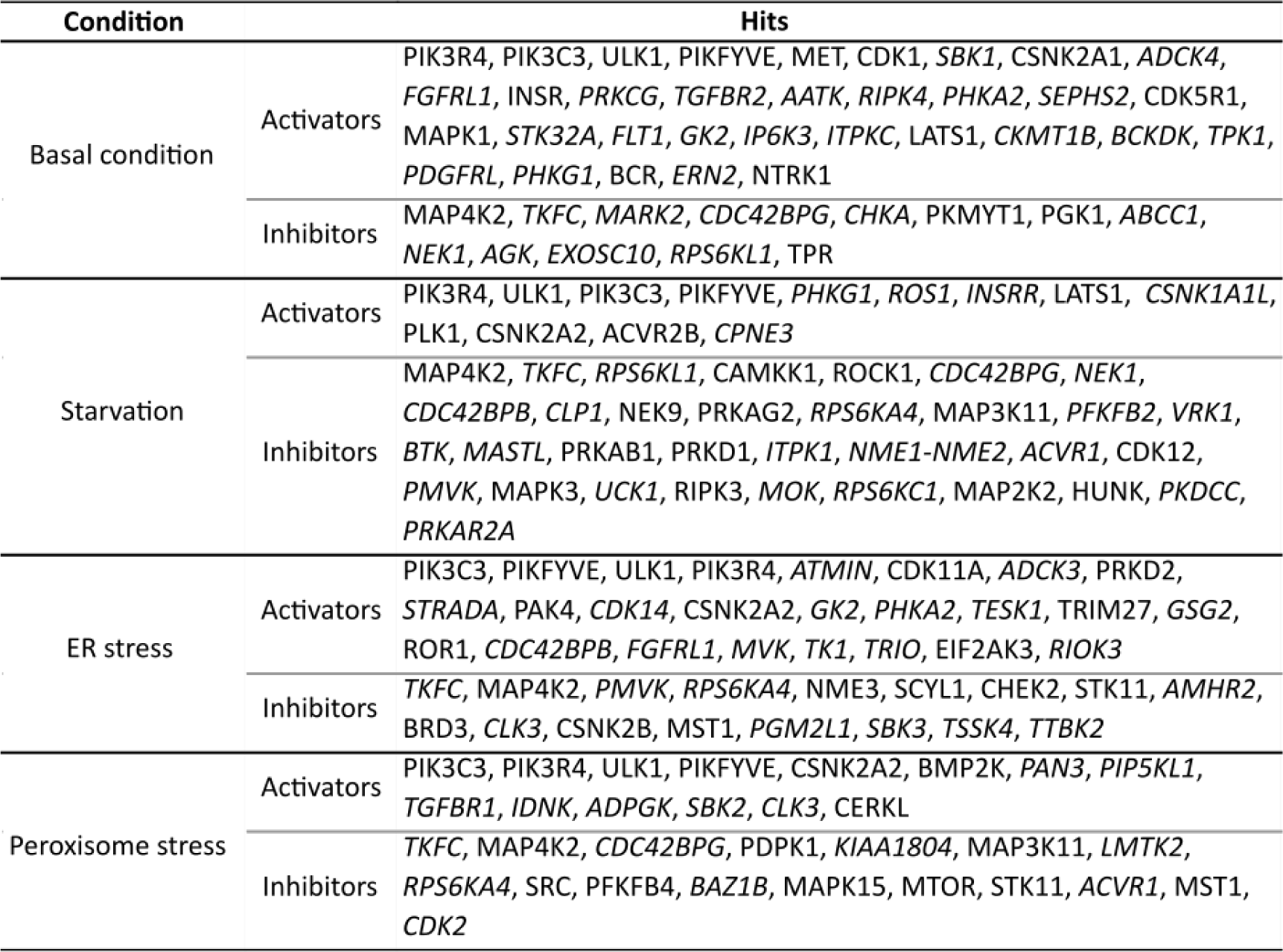
Summary of hits satisfy log_2_FC and FDR cutoffs (known hits are displayed in normal texts and novel hits are shown in italic texts).

#### Autophagy activators

Our kinome screens identified a total of 66 autophagy activators. Notably, the autophagy activators that were found in all four conditions have all previously been linked to autophagy activation, reinforcing the quality of our screen results. The common activators were *ULK1*, *PIK3C3*, *PIK3R4*, and *PIKFYVE* (Figure 2C). The protein kinase ULK1 along with the lipid kinase containing PIK3R4 and PIK3C3 are important factors in core autophagy machinery and have been well characterized to tightly regulate autophagy initiation. Although the role of PIKFYVE in autophagy has only recently been discovered, it is reported to be important for lysosomal function in the autophagy pathway, which is a common requirement for bulk and selective autophagy.

We also found multiple genes encoding members of the casein kinase family (CSNK) among the four autophagy screens. Several reports have demonstrated both activating and inhibiting roles of this family in autophagy regulation, with the role varying based on the cell type and stress condition tested^88,89,90,91,92^. In our analysis, we found *CSNK2A1* was involved in autophagy activation under basal conditions, while *CSNK1A1L* and *CSNK2A2* were implicated in activating starvation-induced autophagy. *CSNK2A2* was also associated with autophagy activation under ER stress and peroxisomal stress (Table 1).

##### Autophagy under basal conditions

We identified 33 basal autophagy activators. Of 33 candidates, 13 have been previously linked to autophagy (Table 1 plain face text) and 20 were uniquely identified in our screens (Table 1 italic text). Of the 33 hits 24 were only identified in the basal autophagy conditions, 4 were common to all conditions, 2 (*LATS1* and *PHKG1*) were shared between basal and starvation-induced autophagy, and 3 (*FGFRL1*, *PHKA2*, and *GK2*) were shared between basal autophagy and ER stress conditions (Table 1, Figure 2B).

##### Starvation-induced autophagy

We found a total of 13 candidates involved in autophagy induction. Of these hits, 8 have been previously reported to modulate autophagy (Table 1 plain face text) and 5 were unique to our screens (Table 1 italic text). 6 of the 13 hits were identified as autophagy activators only under starvation (Table 1, Figure 2B). *LATS1* and *PHKG1* were shared between starvation-induced and basal autophagy Finally, *CSNK2A2* was identified as a shared autophagy activator under starvation, ER stress, and peroxisomal stress.

##### ER stress-induced autophagy

We identified 25 (11 established and 14 novel) kinases associated with autophagy under ER stress (Table 1). Of the 25 hits, 17 were identified only in the ER stress-induced autophagy (Figure 2B), 4 were common to all conditions, 3 (*FGFRL1*, *PHKA2*, and *GK2*) were shared between ER stress conditions and basal autophagy, and 1 (*CSNK2A2*) was found as a shared autophagy activator under starvation, ER stress, and peroxisomal stress (Figure 2B). Notably, our screens identified *EIF2AK3* as a unique ER stress-induced autophagy activator. EIF2AK3, also known as PERK (protein kinase R-like endoplasmic reticulum kinase), is a key player in the unfolded protein response and initiates the upstream response to ER stress by activating transcription factors responsible for expressions of autophagy proteins^93^. Moreover, it was reported that EIF2AK3 is selectively required for ER-phagy induced by tunicamycin^94^, which validates the ability of our screens to recognize selective autophagy modulators.

##### Peroxisomal stress-induced autophagy

We identified 14 candidates (7 established and 7 novel) involved in autophagy induction in response to peroxisomal stress (Table 1). Previously, ATM had been the sole kinase reported to induce pexophagy^54^. We did not identify *ATM* in our screens, which is likely due to the genetic intolerance of HEK293A cells to *ATM* depletion. However, our screens found 9 unique candidates linked to peroxisomal stress-induced autophagy (Table 1). *CSNK2A2* is the only common hit found in peroxisomal stress-induced autophagy and in other autophagy pathways (starvation- and ER stress-induced autophagy).

#### Autophagy inhibitors

We identified a total of 63 shared and distinct autophagy inhibitors across the four screens. We found that *TKFC* and *MAP4K2* are shared autophagy inhibitors among all conditions (Figure 2D, Table 1). TKFC (triokinase/FMN cyclase) is enzyme involved in cellular processes related to the metabolism of carbohydrates and flavin mononucleotide (FMN), which is a type of flavin coenzyme^95^. There is no known link between TKFC and autophagy to date. MAP4K2, mitogen-activated protein kinase kinase kinase kinase 2, is a serine/threonine kinase essential for innate immune responses and cell signaling^96^. Little is known about the relationship between MAP4K2 and autophagy. However, MAP4K2 has been recently reported to induce autophagy upon energy stress through LC3 phosphorylation. *MAP4K2* inclusion in all four investigated conditions underscores its role as a general inhibitor of autophagy^97^.

Our screens also identified genes encoding members of ribosomal protein S6 kinase (RPS6K) family across four tested conditions. Similar to casein kinases, various reports have indicated that this family plays both activating and inhibitory roles in the regulation of autophagy^98,99,100,101,102^. In this study, we only observed *RPS6K* presence in the negative autophagy regulator screens. Specifically, *RPS6KL1* was found in the basal condition. *RPS6KA4*, *RPS6KC1*, and *RPS6KL1* were enriched in starvation-induced autophagy conditions. Finally, *RPS6KA4* was enriched in both ER stress- and peroxisomal stress-induced autophagy.

##### Autophagy under basal conditions

We identified 13 candidates (4 established, 9 novel) involved in basal autophagy (Table 1). 8 of the 13 candidates were identified as autophagy inhibitors only under basal conditions (Figure 2B). *NEK1* and *RPS6KL1* are two hits involved in basal and starvation-induced autophagy. *CDC42BPG* is the sole candidate shared among basal, starvation-induced, and peroxisomal stress-induced autophagy processes.

##### Starvation-induced autophagy

Our kinome screens found a total of 33 candidates (13 established, 20 novel) associated with autophagy inhibition in response to starvation (Table 1). 24 of these were identified only in the starvation conditions, 2 were common to all conditions, 2 (*NEK1* and *RPS6KL1*) were shared between starvation-induced and basal autophagy, 1 (*PMVK*) was found in autophagy inhibition under starvation and ER stress, 2 (*MAP3K11* and *ACVR1*) were identified in starvation- and peroxisomal stress-induced autophagy, 1 (*CDC42BPG*) was shared among basal, starvation-induced, and peroxisomal stress-induced autophagy processes, and 1 (*RPS6KA4*) was found in starvation-, ER stress-, and peroxisomal stress-induced autophagy.

##### ER stress-induced autophagy

We identified 17 hits (8 established, 9 novel) involved in autophagy inhibition under ER stress (Table 1). We found 11 hits selectively associated with ER-phagy (Figure 2B). Notably, CSNK2B is the only member within the casein kinase family that we identified in the context of autophagy inhibition. In addition to the overlapped hits mentioned above, *STK11* and *MST1* are candidates shared among ER stress- and peroxisomal stress-induced autophagy.

##### Peroxisomal stress-induced autophagy

Our screens found 17 candidates (9 established, 8 novel) associated with autophagy inhibition in response to peroxisomal stress (Table 1). 9 of these were found only in the peroxisomal stress conditions, 2 were common to all conditions, 2 (*MAP3K11* and *ACVR1*) were identified in peroxisomal stress- and starvation-induced autophagy, 2 (*STK11* and *MST1*) were shared among peroxisomal stress- and ER stress-induced autophagy, 1 (*CDC42BPG*) was found in basal, starvation-, and peroxisomal stress-induced autophagy processes, and 1 (*RPS6KA4*) was involved in autophagy inhibition under peroxisomal stress, starvation, and ER stress.

Collectively, our comprehensive kinome screens have identified a total of 129 kinases with a significant FDR and positive log_2_FC values. In addition to the common activators and inhibitors of all conditions, we also found several kinases that overlapped between 2 or more conditions, which may imply common upstream regulation between these types of autophagy. There is a precedent for this type of overlap in selective autophagy regulators from the study of autophagy receptors. For example, BNIP3L has been reported to induce pexophagy and mitophagy^103^. In addition to shared regulators, we found selective regulators of both pexophagy and ER-phagy, indicating that selective autophagy is likely governed by signal transduction in addition to autophagy receptors.

### Selection of top hits for characterization

To validate the hits as regulators of selective autophagy regulation, we generated polyclonal knockout (KO) cell lines using CRISPR/Cas9 for the top activators or inhibitors for both ER stress-induced and peroxisomal stress-induced autophagy and validated them in an orthogonal system. An additional ULK1 positive control, which was shared among all conditions, was also generated in the same manner. We evaluated p62 flux via WB following the same treatment paradigms used in the screens (Figures S2 and S3). From this experiment, we chose the top hits for each category (ER-phagy/pexophagy activator/inhibitor), which showed the largest average regulatory effect and had not previously been implicated in regulation of the corresponding selective autophagy pathway. These hits were CDK11A (ER-phagy activator), NME1 (ER-phagy repressor), PAN3 (pexophagy activator), and CDC42BPG (pexophagy inhibitor); KO efficiencies of these cell lines were confirmed by WB (Figure S4).

### CDK11A is a selective ER-phagy activator

Our analysis suggested that CDK11A may be a novel activator of ER-phagy (Figures 3A and S2A). CDK11A (Cyclin-Dependent Kinase 11A), a member of the cyclin-dependent kinase (CDK) family, is implicated in various cellular processes, including regulation of transcription, cell cycle control, and basal autophagy regulation. However, its role in selective autophagy process is unclear. We treated control 293A, CDK11A KO, or ULK1 KO cells with tunicamycin, and observed a decrease of p62 in wildtype cells, but not in either ULK1 KO or CDK11A KO cells (Figure 3B). Moreover, we observed an increase in basal p62 in both ULK1 KO and CDK11A KO cells, which is expected when an activator of autophagy is ablated (Figure 3B)^104^. Tunicamycin is known to induce the expression of genes in the unfolded protein response including C/EBP-homologous protein (CHOP)^105^. Western blot analysis of CHOP showed an equivalent activation of the UPR pathway in all 3 cell lines upon tunicamycin treatment, indicating that an altered unfolded protein response was not responsible for differences in p62 flux (Figure 3B).

**Figure 3:**
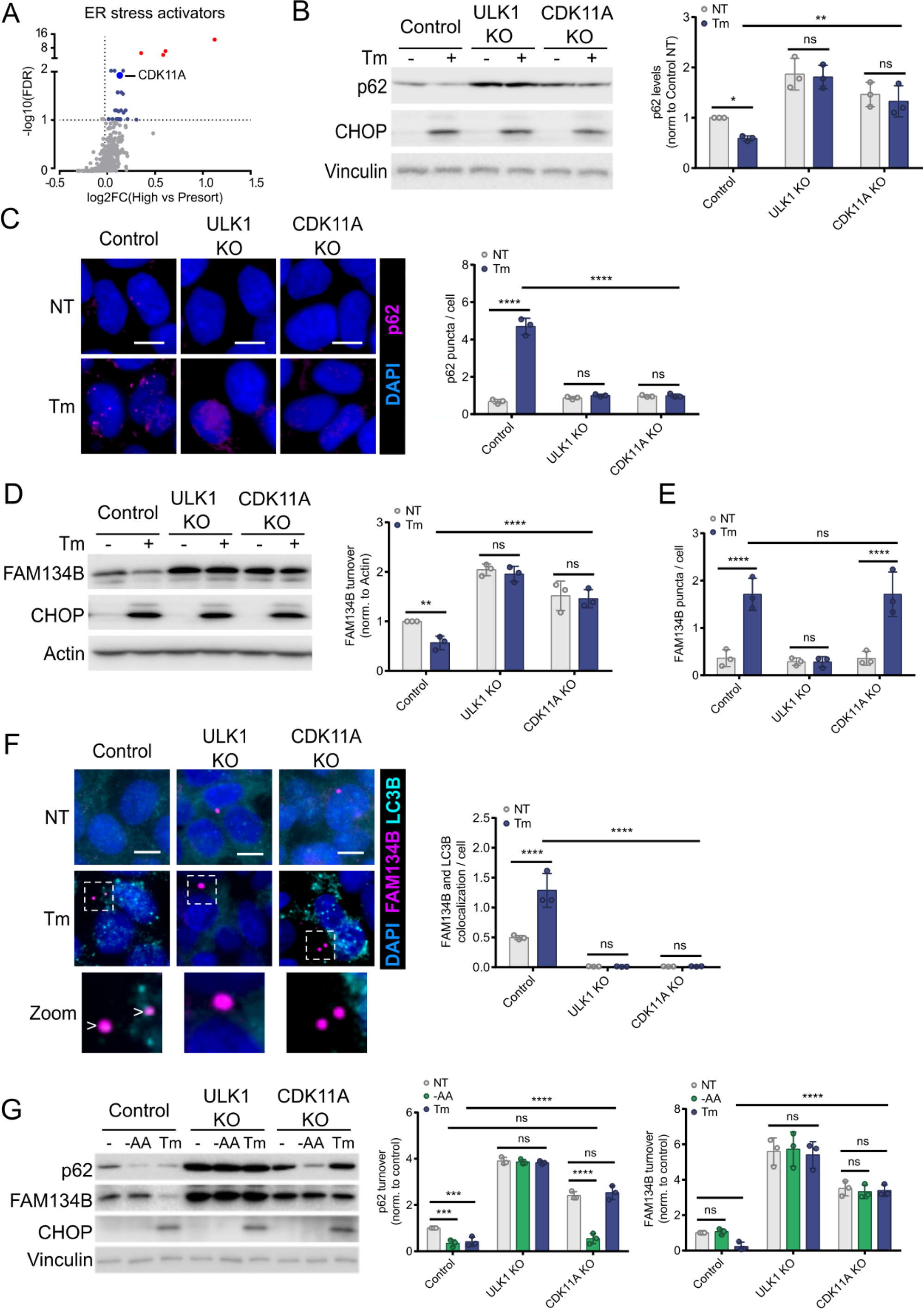
CDK11A activates ER-phagy. (A) CDK11A is represented as a prominent blue dot on the volcano plot. Red dots on the graph denote common regulators across all four conditions. (B) The control, ULK1 KO, and CDK11A KO cells were treated with tunicamycin (10 µg/mL) for 6 hours. Levels of p62 were analysed using western blot. The effectiveness of tunicamycin was assessed through CHOP analysis. (C) The control and KO cells were incubated with tunicamycin (10 µg/mL) for 6 hours. p62 puncta were visualized and quantified by immunofluorescence. Scale bars, 10 µM. (D) The control and KO cells were treated with tunicamycin. FAM134B levels were then examined using western blot. (E) The indicated cells were incubated with tunicamycin for 6 hours. FAM134B puncta were quantified by immunofluorescence. (F) The control and KO cells were incubated with tunicamycin for 6 hours. FAM134B and LC3B puncta were visualized and quantified by immunofluorescence. White arrows depict LC3B and FAM134B colocalization. Scale bars, 10 µM. (G) The control and KO cells were incubated with either amino acid-free media (1.5 hours) or tunicamycin (10 µg/mL, 6 hours). Whole-cell lysates were immunoblotted using the antibodies indicated. Unless otherwise indicated, experiments were performed three times. Data are represented as means ± SDs, and p values were determined by two-way ANOVA. *p□≤□0.1; **p□≤□0.01; ***p□≤□0.001; ****p□≤□0.0001; ns, not significant.

We used immunofluorescence (IF) microscopy as an orthogonal approach to determine whether CDK11A regulates p62 during ER stress. In response to ER stress, wild-type cells show an increase in p62 puncta, which is indicative of p62 loading in autophagosomes (Figure 3C). However, ULK1 KO and CDK11A KO cells failed to mount this response, further supporting the notion that both genes are required for autophagy induction in response to ER stress (Figure 3C).

To better link CDK11A function to ER-phagy induction, we turned to the ER-resident autophagy receptor: FAM134B. Among the ER-phagy receptors, the membrane embedded FAM134B is the best characterized^45,106^. Like p62, FAM134B it is loaded into autophagosomes and degraded. However, unlike p62, FAM134B is specifically linked to ER-phagy. We treated wild-type, CDK11A KO, or ULK1 KO cells with tunicamycin and analysed FAM134B levels. We observed a decrease of FAM134B in wildtype cells, but not in either ULK1 KO or CDK11A KO cells, confirming that stress-induced ER-Phagy requires CDK11A (Figure 3D). We next immunostained for FAM134B to determine if FAM134B loading into autophagosomes upon ER stress was inhibited by CDK11A KO. Interestingly, we observed that FAM134B puncta were elevated in both tunicamycin-treated wildtype CDK11A KO cells (Figure 3E). FAM134B interacts with LC3B through LIR motifs, which guides the sequestration and engulfment of ER fragments within autophagosomes^45^. To distinguish if FAM134B puncta were associated with functional autophagosomes, we repeated the experiment while co-staining for LC3B. As expected, control cells showed an induction of LC3B-associated FAM134B puncta under ER stress conditions, indicating an activation of ER-phagy (Figure 3F). However, in CDK11A or ULK1 deficient conditions, we could not detect colocalization between FAM134B and LC3B puncta, likely indicating stalled autophagic structures. This is consistent with our WB analysis (Figure 3D) and previous reports of stalled autophagic structures in autophagy-deficient cells^104,107,108^. Interestingly, ER stress induced LC3-negative FAM134B puncta in CDK11A KO cells, but not ULK1 KO cells (Figure 3F), indicating that ULK1 may be required for FAM134B oligomerization, while CDK11A regulates ER-phagy at a step downstream of FAM134B oligomerization, but upstream of LC3B binding. Collectively, these data indicate that CDK11A is an essential activator of ER-phagy upon ER stress.

To determine the selectivity of CDK11A towards ER-phagy, we tested how its loss of function affects starvation-induced autophagy. To make this head-to-head comparison, we incubated wildtype, ULK1 KO, or CDK11A KO cells with amino acid-free media or tunicamycin. As expected, we observed an efficient induction of starvation-induced autophagy and ER-phagy in wildtype cells as shown by the degradation of p62 and FAM134B, respectively (Figure 3G). While ULK1 KO cells showed an expected deficiency in both starvation-induced autophagy and ER-phagy, we observed that CDK11A KO cells were competent in the induction of starvation-induced autophagy, but deficient for ER-phagy (Figure 3G). Thus, our data identify CDK11A as a selective activator of ER-phagy.

### NME3 is a selective ER-phagy inhibitor

Nucleoside Diphosphate (NDP) Kinase 3 (NME3) is best known for its ability to regulate nucleotide metabolism and signaling^109,110^. Recent research has indicated that NME3 serves two distinct functions involving regulation of mitochondrial dynamics and NDP kinase activity, both of which are required to maintain cell viability under glucose starvation^111^. Additionally, during the writing of this manuscript, NME3 was been reported to be involved in mitophagy regulation^112^. Our analysis of the top screen hits indicated that NME3 may be a selective repressor of ER-phagy (Figures 4A and S2B). We first tested the ability of NME3 KO cells to increase autophagic flux upon ER stress. Compared to our wildtype cells, we found NME3 KO cells exhibited lower levels of p62 and a more complete degradation of p62 under ER stress, consistent with a role for NME3 as an inhibitor of ER-phagy (Figure 4B). Next, we analysed the impact of NME3 KO on ER stress-induced p62 loading into autophagosomes. As expected, we observed an increase in p62 puncta in the treated control cells and a lack of p62 signal in both treated and untreated ULK1 KO cells (Figure 4C). Conversely, we found that NME3 KO cells treated with tunicamycin showed more p62 puncta compared to those of the treated control cells (Figure 4C). Together, these data show that NME3 regulates autophagy flux in response to ER stress. We then examined whether NME3 could regulate ER-phagy receptor, FAM134B. The control and KO cells were exposed to tunicamycin and FAM134B levels were investigated using WB. Consistently, NME3 depletion resulted in a reduction in FAM134B levels compared to those of the control cells upon ER stress (Figure 4D). The changes in FAM134B levels were also quantified (Figure 4D, right panel). Together, these experiments indicate that NME3 regulates ER-phagy.

**Figure 4:**
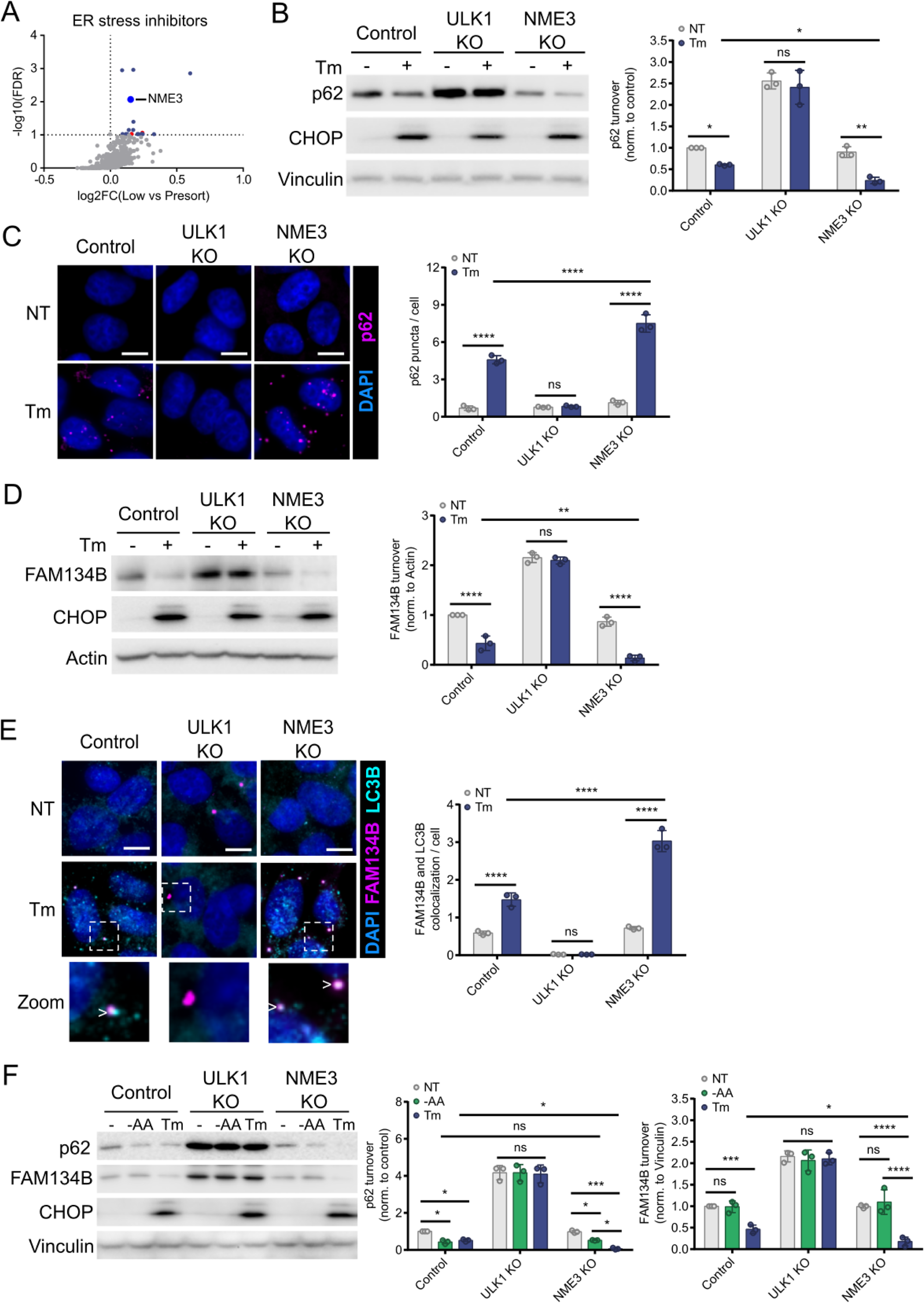
NME3 inhibits ER-phagy. (A) NME3 is represented as a prominent blue dot on the volcano plot. Red dots on the graph denote common regulators across all four conditions. (B) The control, ULK1 KO, and NME3 KO cells were treated with tunicamycin (10 µg/mL) for 6 hours. Changes in p62 levels were analysed using western blot. The effectiveness of tunicamycin was assessed through CHOP analysis. (C) The control and KO cells were incubated with tunicamycin (10 µg/mL) for 6 hours. p62 puncta were visualized and quantified by immunofluorescence. Scale bars, 10 µM. (D) The control and KO cells were treated with tunicamycin. FAM134B signaling was then examined using western blot. (E) The indicated cells were incubated with tunicamycin. FAM134B and LC3B puncta were visualized and quantified by immunofluorescence. White arrows depict LC3B and FAM134B colocalization. Scale bars, 10 µM. (F) The control and KO cells were incubated with either amino acid-free media (1.5 hours) or tunicamycin (10 µg/mL, 6 hours). Whole-cell lysates were immunoblotted using the antibodies indicated. Unless otherwise indicated, experiments were performed three times. Data are represented as means ± SDs, and p values were determined by two-way ANOVA. *p□≤□0.1; **p□≤□0.01; ***p□≤□0.001; ****p□≤□0.0001; ns, not significant.

Next, we investigated the effects of NME3 depletion on FAM134B recruitment into autophagosomes in response to ER stress. We immunostained FAM134B and LC3B in our untreated and treated wildtype or KO cells. We observed NME3 KO cells had an enhanced production of LC3B-positive FAM134B puncta upon ER stress compared to wildtype cells (Figure 4E). The increase of autophagosome-associated FAM134B under stress is a strong indicator that NME3 inhibits ER-phagy rates. We next examined whether NME3 could regulate ER-phagy selectively by treating the wildtype and KO cells with either amino acid-free media or tunicamycin. Consistent with the results above in the NME3 KO background, we observed an increase in p62 and FAM134B degradation upon tunicamycin treatment (Figure 4F). However, NME3 KO cells showed no detectable difference in p62 clearance under starvation when compared to wildtype cells (Figure 4F). Taken together, these experiments show that NME3 is a selective upstream inhibitor of ER-phagy.

### PAN3 is an activator of pexophagy

Our screen validation identified Poly(A)-specific ribonuclease subunit 3 (PAN3) as a potential regulator of clofibrate-driven pexophagy (Figures 5A and S3A). PAN3 plays a crucial role in mRNA degradation and regulation of gene expression in eukaryotic cells^113^. It is a component of the PAN2-PAN3 de-adenylation complex responsible for shortening the poly(A) tail of mRNA molecules^114,115^. Pan3 has a PKc kinase domain but may be a pseudokinase and has no known targets^116^. To test if PAN3 regulates peroxisomal stress-induced autophagy we first exposed the wildtype, ULK1 KO, and PAN3 KO cells to clofibrate (1 mM) for 6 hours. p62 levels were decreased in the wildtype cells, but not in ULK1 KO cells (Figure 5B). Interestingly, p62 flux in response to clofibrate was also blocked in PAN3 KO cells (Figure 5B), suggesting that PAN3 plays a role in modulating autophagy under peroxisomal stress. We next inspected changes in p62 puncta formation in response to clofibrate in wildtype and KO cells. We observed an increase in p62 puncta in response to clofibrate in the wildtype, but not ULK1 KO cells (Figure 5C). However, PAN3 KO cells exhibited a low level of p62 puncta under basal conditions that, unlike the wildtype cells, did not change in response to clofibrate (Figure 5C). Together, these data indicate that PAN3 is required for autophagic induction in response to peroxisomal damage.

**Figure 5:**
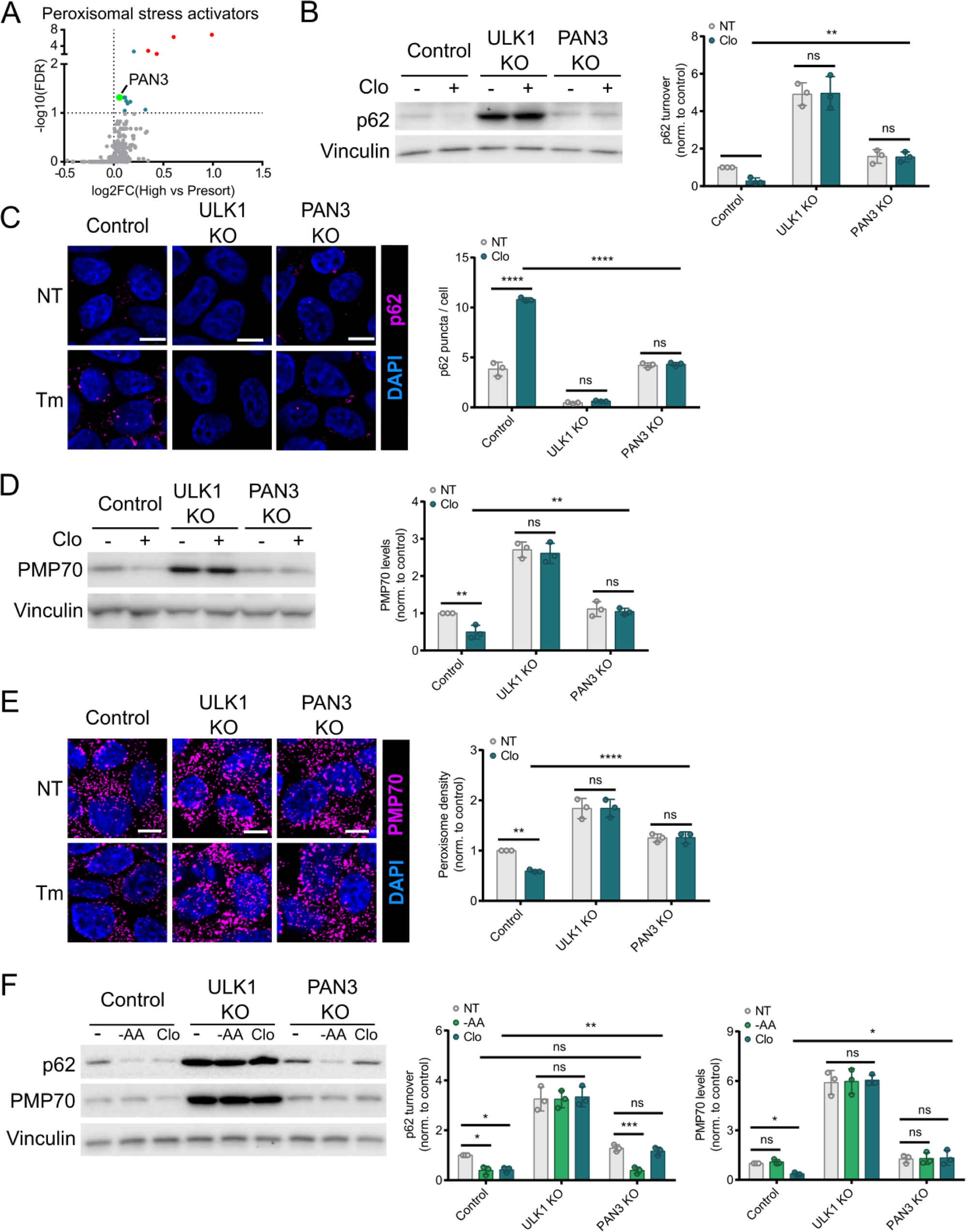
PAN3 activates pexophagy. (A) PAN3 is represented as a prominent teal dot on the volcano plot. Red dots on the graph denote common regulators across all four conditions. (B) The control, ULK1 KO, and PAN3 KO cells were treated with clofibrate (1 mM) for 6 hours. Changes in p62 levels were analysed using western blot. (C), (E) The indicated cells were incubated with clofibrate. Either p62 (C) or PMP70 (E) signal was visualized and quantified by immunofluorescence. Scale bars, 10 µM. (D) The control and KO cells were treated with clofibrate. Western blot was then used to examine pexophagy receptor, PMP70. (F) The control and KO cells were incubated with either amino acid-free media (1.5 hours) or clofibrate (1 mM, 6 hours). Whole-cell lysates were immunoblotted using the antibodies indicated. Unless otherwise indicated, experiments were performed three times. Data are represented as means ± SDs, and p values were determined by two-way ANOVA. *p□≤□0.1; **p□≤□0.01; ***p□≤□0.001; ****p□≤□0.0001; ns, not significant.

Next, to determine the impact of PAN3 on peroxisome turnover, we used an established peroxisomal marker, peroxisomal membrane protein 70 (PMP70). Upon stress, PMP70 is ubiquitinated by peroxisomal E3 Ub ligase PEX2 and promotes pexophagy, thus maintaining peroxisome quality^53^. Consistent with previous reports, we found that in wildtype cells clofibrate caused a reduction in PMP70 indicating a reduction in peroxisomes. This peroxisomal loss was dependent on autophagy as this decrease was not observed in clofibrate-treated ULK1 KO cells (Figure 5D). Interestingly, clofibrate-treated PAN3 KO cells exhibited normal PMP70 levels, which is indicative of a defect in pexophagy activation (Figure 5D). To confirm this observation, we next examined peroxisome density in the untreated and treated wildtype and KO cells through PMP70 staining. We observed a decrease in the PMP70 signal in the clofibrate-treated wildtype cells, with an elevated and clofibrate-insensitive population of peroxisomes in ULK1 KO cells (Figure 5E). Importantly, in PAN3 KO cells we observed no loss of peroxisome density in response to clofibrate (Figure 5E). Collectively, these experiments showed that PAN3 is an activator of pexophagy, which is required for clearance of peroxisomes damaged by clofibrate.

We next sought to examine the ability of PAN3 to regulate specific pexophagy. In addition to clofibrate treatment, we incubated the control and KO cells with starvation media for a short period of time, during which PMP70 levels were not affected (Figure 5F). We then investigated the changes in p62 levels. While PAN3 KO cells exhibited a defect in clofibrate-stimulated pexophagy, they exhibited a normal induction of starvation-induced autophagy (Figure 5G). Taken together, these data demonstrate that PAN3 is a selective activator of pexophagy.

### CDC42BPG is an inhibitor of pexophagy

CDC42-binding protein kinase gamma (CDC42BPG) is a serine/threonine protein kinase known to interact with the small GTPase CDC42 and is involved in cytoskeletal organization, cell division, and cell migration^117^. To date CDC42BPG has not been linked to autophagy regulation or peroxisomes. Our kinome screen and subsequent analysis revealed involvement of CDC42BPG in the suppression of autophagy triggered by peroxisomal stress (Figures 6A and S3B). Notably, CDC42BPG is the shared candidate inhibiting autophagy under basal condition, starvation, and peroxisomal stress. We decided to further validate CDC42BPG as it appears as a top novel hit displaying highly significant FDR and log_2_FC values and its depletion showed the most robust response upon clofibrate treatment (Figure S3B). Indeed, when we exposed wildtype, ULK1 KO, and CDC42BPG KO cells to clofibrate, we observed that the reduction of p62 in CDC42BPG KO cells was significantly higher than in the wildtype cells, indicating that it may inhibit autophagy induced by peroxisomal stress (Figure 6B). Additionally, p62 levels were not significantly changed under basal conditions in wildtype and CDC42BPG KO cells, suggesting that it is dispensable for basal autophagy (Figure 6B). We further examined clofibrate-induced p62 puncta formation by IF. In line with previous observations, there was an upregulation of p62 puncta in the clofibrate-incubated control cells, while there was a lack of p62 signals in both unstimulated and stimulated ULK1 KO cells (Figure 6C). Notably, we observed a more significant increase in p62 puncta in the treated CDC42BPG compared to those of the treated control cells (Figure 6C). Collectively, these data show that CDC42BPG plays a role in inhibiting clofibrate-induced autophagic flux.

**Figure 6:**
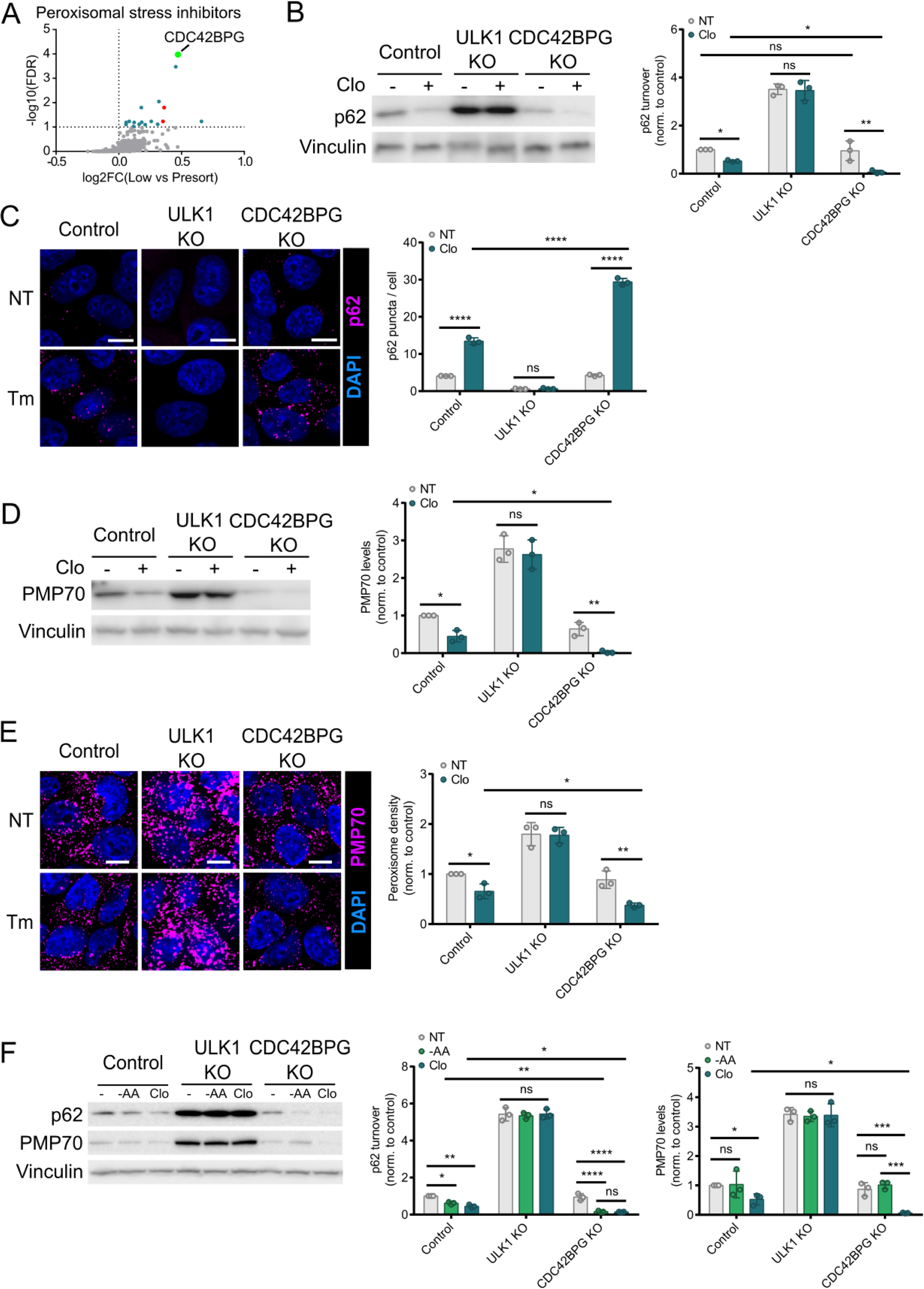
CDC42BPG inhibits pexophagy. (A) CDC42BPG is represented as a prominent teal dot on the volcano plot. Red dots on the graph denote common regulators across all four conditions. (B) The control, ULK1 KO, and CDC42BPG KO cells were treated with clofibrate (1 mM) for 6 hours. Changes in p62 levels were analysed using western blot. (C), (E) The indicated cells were incubated with clofibrate. Either p62 (C) or PMP70 (E) signal was visualized and quantified by immunofluorescence. Scale bars, 10 µM. (D) The control and KO cells were treated with clofibrate. PMP70 was then analysed using western blot. (F) The control and KO cells were incubated with either amino acid-free media (1.5 hours) or clofibrate (1 mM, 6 hours). Whole-cell lysates were immunoblotted using the antibodies indicated. Unless otherwise indicated, experiments were performed three times. Data are represented as means ± SDs, and p values were determined by two-way ANOVA. *p□≤□0.1; **p□≤□0.01; ***p□≤□0.001; ****p□≤□0.0001; ns, not significant.

Next, we investigated the effects of CDC42BPG on PMP70 protein levels. We observed a more pronounced downregulation in PMP70 levels in the clofibrate-treated CDC42BPG KO cells compared to those of stressed wildtype cells (Figure 6D). We next investigated stress-dependent changes in peroxisome density by IF in the wildtype and KO cells through PMP70 staining. We saw a decrease in peroxisome density in the clofibrate-stressed wildtype cells, which was absent in ULK1 KO cells (Figure 6E). Notably, we observed a more robust stress-induced clearance of peroxisome staining in the CDC42BPG KO cells, compared to wildtype control cells (Figure 6E). Collectively, these experiments demonstrate that CDC42BPG is a negative regulator of pexophagy.

We next asked whether CDC42BPG could modulate pexophagy specifically by exposing the control and KO cells with either amino acid-free media or clofibrate. As expected, we saw stress-induced p62 downregulations in the treated wildtype cells, which was absent in ULK1 KO cells (Figure 6F). Notably, starvation-induced p62 degradation in CDC42BPG KO cells was slightly higher than that in wildtype cells (Figure 6F). This was not unexpected as CDC42BPG also appeared as a negative regulator in the starvation condition. In summary, our findings suggest that CDC42BPG is capable of inhibiting autophagy, with its most robust inhibition impacting pexophagy.

## Discussion

In this study, we have used a pooled CRISPR screening workflow to systematically identify both activators and inhibitors of autophagy under multiple conditions, including basal state, starvation, ER stress, and peroxisomal stress. Some of these regulators were common to selective and bulk autophagy, and have been previously linked to autophagy. These include PIK3C3 (VPS34), PIKFYVE, PIK3R4 (VPS15), and ULK1. In addition, we identified shared autophagy inhibitors that are not currently described to supress autophagy including MAP4K2 and TKFC. TKFC is a member of the dihydroxyacetone kinase family, which is best described to phosphorylate glyceraldehyde to glyceraldehyde-3-phosphate^118^. It will be interesting to determine if the metabolic impacts of TKFC knockout are responsible for autophagy activation, or if the autophagy regulation is through an alternate function of TKFC. Notably, TKFC phosphorylation of glyceraldehyde is utilized in the metabolism of fructose, which is not present in our culture media, highlighting the possibility of an alternate mechanism of regulation^118^. Interestingly, MAP4K2 came up as an inhibitor in our screen, but has been described in one study as an autophagy activator^97^. However, there were notable differences in the studies, which might explain the differences between our observations. For example, MAP4K2 activated autophagy under 20-hour glucose starvation, while we identified MAP4K2 as an autophagy inhibitor under 3-hour amino acid starvation. Chronological dissection of nutrient deprivation will help tease out its potential dual role in autophagy regulation. It will also be interesting to test whether MAPK4K2-linked inhibition is mediated by its kinase activity towards LC3, which may be functionally impacted by competing or proximal phosphorylation by PKA or PKCλ^119^. These open questions raised from the hits described above indicate that further characterization of our hits for regulators of bulk autophagy may also provide insight into basal and starvation-induced autophagy. While not the focus of our screens, we identified: 25 potential novel regulators of starvation-induced autophagy, and 29 potential novel regulators of basal autophagy.

In response to ER stress our screen identified an enrichment of 25 kinases linked to autophagy activation and 17 with inhibition. To validate the specificity of these regulators we characterized CDK11A and NME3 as selective regulators of ER-phagy. Specifically, we found CDK11A was a selective activator of ER-phagy and not required for starvation-induced autophagy. CDK11 has established role in control of RNA splicing, transcription, and the cell cycle control. Interestingly, dual siRNA-mediated knockdown of CDK11A and CDK11B has previously been linked to an acute activation of basal autophagy, followed by an inhibition of autophagy at a later time point^120^. However, the mechanism of regulation remains unknown. In our knockout cells disruption of CDK11A was sufficient to significantly impair ER-phagy induction, without any detectible enrichment in basal or starvation conditions. However, it remains to be seen if a dual knockout of CDK11A and CDK11B would impact basal autophagy, or if the basal autophagy phenotype previously observed may involve a defect in ER homeostasis. CDK11A and CDK11B are activated in multiple cancer types and have been linked to acquisition of oncogenic properties, including proliferation. This newly described function of CDK11A in ER-phagy begs the question of whether ER stress dysregulation may be integral to these oncogenic properties.

We found that NME3 is a selective repressor of ER-phagy. NME3 belongs to a more conserved group of nucleoside diphosphate kinase (NDPK) family, which regulates cellular nucleotide homeostasis and is associated with GTP-dependent cellular processes^121^. However, independent of its NDPK activity, NME3 has been described to regulate mitochondrial dynamics^122^. Both NDPK and mitochondrial functions are important for cellular survival under glucose starvation^122^. Moreover, it has been recently reported that NME3 is important for mitophagy induction^112^. It will be interesting in future studies to determine whether NME3-mediated repression of ER-phagy is linked to its involvement in mitophagy and to elucidate the mechanisms by which NME3 performs conflicting roles in selective autophagy. An inactivating autosomal recessive mutation NME3 was found in a case study of rare consanguineous fatal neurodegenerative disorder^122^. While homozygous inactivating mutations are lethal early in life, the impact of heterozygous inactivation of NME3 on ER-phagy and any potential physiological consequences is an interesting area for investigation.

In response to peroxisomal stress we identified an enrichment of 14 autophagy activators and 17 inhibitors. To validate the specificity of these regulators we chose to characterize the activator PAN3 and inhibitor CDC42BPG. PAN3 is a component of PAN2-PAN3 complex, which modulates mRNA stability or translational efficiency and has not been implicated in autophagy^116^. Additional work is required to determine if PAN3 regulates gene expression of pexophagy promoters, or whether its promotion of peroxisomal autophagy is mediated by an alternate mechanism. CDC42BPG is a less well characterized member of the Myotonic dystrophy-related Cdc42-binding kinases (MRCK), which play an important role in actin-myosin regulation and other functions such as cell invasion, motility, and adhesion^117^. While neither of these genes linked to disease, the mechanisms of peroxisomal disorders such as Zellweger’s disease have not been fully elucidated and exhibit dysregulation of pexophagy. Therefore, it would be interesting to test the involvement of hits from our pexophagy screen, including these proteins in cases which do not have a reported Zellweger-associated PEX mutations.

Our screen design offers several advantages. First, the employment of CRISPR/Cas9 system provides higher efficiency and lower off-target effects compare to traditional RNAi screens. Permanent gene perturbation provides more robust signals. The screen design allows cells to have more time to recover and have more optimal responses to stress conditions. Utilization of p62 reporter enables investigation of different selective autophagy processes owing to its ability to bind diverse ubiquitinated substrates. Finally, fixation of samples prior to sorting allows for concurrent analysis of different stress-induced autophagy pathways. Although this screening approach has clear benefits, it also possesses limitations. This CRISPR/Cas9 system typically results in complete loss of function of genes, making it impractical for studying essential genes or partial loss of functions. Adaptations of RNAi or CRISPRi to this read-out can help complement these findings. Another limitation of this study is the choice of reporter. While p62 is linked to several types of selective autophagy, the relative clearance of total p62 under each condition may be different depending on cargo abundance and preference for other autophagy receptors. As such, the detection threshold was likely different among the conditions tested. Organelle-specific autophagy receptors would be ideal for sensitivity, with the trade-off of losing the ability to directly compare each stress condition.

Together, these screens have identified a heretofore underappreciated role for signal transduction pathways in the regulation of selective autophagic pathway. This resource thus provides a host of putative regulators, paving the way for tighter, selective control of different forms of autophagy and potential therapeutic inroads to target these pathways in clinically relevant scenarios.

## Material and methods

### Antibodies and reagents

Anti-ULK1 (Cat#6439S, 1:1000) antibody was obtained from Cell Signaling Technology. Anti-LC3B (Cat#PM036 for immunofluorescence, 1:2000) and anti-p62 (Cat#M162-3 for immunofluorescence, 1:400) antibodies were purchased from MBL. Anti-PMP70 (Cat#ab3421 for immunofluorescence, 1:1000) antibody was purchased from Abcam. Anti-beta-actin (Cat#A5441 clone AC-15, 1:30K), anti-vinculin (Cat#V9131, 1:30K), and anti-PMP70 (Cat#SAB4200181 for WB, 1:1000) antibodies were obtained from Sigma. Anti-p62 (Cat#sc-28359, 1:1000), anti-PAN3 (Cat#sc-376434, 1:500), and anti-CDC42BPG (Cat#sc-517148, 1:500) antibodies were obtained from Santa Cruz Biotechnology. Anti-FAM134B (Cat#21537-1-AP, 1:1000), anti-NME3 (Cat#15136-1-AP, 1:500), and anti-CHOP (Cat#15204-1-AP, 1:1000) antibodies was obtained from Proteintech Group. Anti-tRFP (Cat#AB233, 1:1000) was purchased from Evrogen. Anti-CDK11A (Cat#ARP61814_P050, 1:500) was obtained from Aviva Systems Biology.

### Cell culture

HEK293A were cultured in DMEM supplemented with 10% bovine calf serum (VWR Life Science Seradigm). Amino acid starvation media was prepared based on Gibco standard recipe omitting all amino acids and supplemented as above without addition of non-essential amino acids and substitution with dialysed FBS (Invitrogen). Media was changed 24 h before experiments.

### Virus generation and concentration

Lentiviral vectors (LentiCRISPRv2 or pCLIP-dual) and their corresponding packaging vectors (psPAX2 and pMD2G) were co-transfected into HEK293T cells in a 4:3:1 molar ratio, respectively. Media was changed 16 hr following transfection to low volume media (5 mL for a 10 cm dish). Media was collected at 48 hr following transfection, replaced with fresh media (5 mL), and collected again at 72 hr. Viral supernatant was filtered through a 0.45 µM polyethersulfone membrane (VWR). Cleared supernatants were concentrated using Virus Precipitation Kit (Benchmark Bioscience) to 1/100 of the original volume.

### Generation of knock-out cell lines using CRISPR/Cas9

sgRNA pairs targeting genes of interest were selected from the transEDIT-dual CRISPR Whole Genome Arrayed Library (Transomic Technologies, Huntsville, AL). They were used in conjunction with a Cas9 expression vector containing neomycin (G418) resistance transcript (Addgene #98292). H293T cells were transfected with lentivirus packaging plasmids and plasmids carrying either sgRNAs or Cas9. The media was collected 4 times throughout the course of 3 days and was filtered through a 0.45 µm syringe filter. Next, wild-type HEK293A cells were infected with both lentiviruses harboring the Cas9 and sgRNAs. The transduced cells were then selected with puromycin (1 µg/mL; 3 days) followed by G418 (1 mg/mL; 6 days). CDK11A sgRNA sequences (5’➔3’): GATTGTGGTGGGCAGCAACA and GATCGATTTCCGAATTCCCG. NME3 sgRNA sequences: CCGCGGGGATTTCTGCATCG and CTTCGCTAACCTCTTCCCCG. PAN3 sgRNA sequences: GTCTCCAGTCTCTGACCAAG and CCGCCCGCGACGGCTCCCGG. CDC42BPG sgRNA sequences: CCATCGATGTGTTTGACGTG and TCGACTTGCGCTTGGCACCG.

### Generation of stable cell lines

HEK293A cells were transduced with lentiviruses carrying DsRed-IRES-GFP-p62. These cells underwent G418 selection and were sorted into single cell populations. FACS was utilized to identify a monoclonal population expressing optimal GFP:DsRed ratio and responses to known autophagy stimuli. Knockout populations used for screen validation were generated by transducing parental HEK293A with sgRNAs targeting potential hits and Cas9. These cells were subjected to puromycin and G418 selection.

### Flow cytometry

Following defined treatments, cells were fixed with 2% paraformaldehyde (PFA) for 10 min and incubated with Tris (pH 8, direct addition to 2% PFA to create a final concentration of 1 M) for 15 min at room temperature. Media were removed. The cells were then harvested using scrapers, resuspended in ice-cold flow buffer (1% BSA and 2 mM EDTA in PBS), and filtered using cell strainers (70 µM, Falcon). The fixed samples were analysed using a BD FACSCelesta flow cytometer. For sorting, the cells were subjected to FACS on a Sony SH800S cell sorter.

### Pooled kinome-wide CRISPR/Cas9 screens

#### Cell culture

The 293A cells expressing the DsRed_IRES-GFP-p62 transgene were plated at approximately 7.5 million cells on 15-cm plates. Next day, these cells were transduced with lentiviruses carrying human kinome CRISPR knockout pooled library at Multiplicity of Infection of 0.3 (13 million cells were infected with 3.86 million Transduction Units to achieve approximately 1000-fold representation of each sgRNA) in the presence of 10 mg/ml polybrene. The library was purchased from Addgene (Cat#1000000083) and amplified using the protocol provided by Addgene. The transduced cells were then selected with puromycin (1 µg/mL) for 3 consecutive days and cultured for an additional 11 days to allow for effective target knockout^123^. We found that day 16 was the earliest for achieving an optimal autophagic response following knockout and recovery from selection. Thus, on day 16, the cells were treated with the stress conditions described above and fixed with 2% PFA for 10 min at room temperature, followed by 15 min incubation with Tris (pH 8, direct addition to 2% PFA to create a final concentration of 1 M). Following aspiration of media, the cells were collected using scrapers, stored in ice-cold flow buffer (1% BSA and 2 mM EDTA in PBS), and filtered using cell strainers (70 µM, Falcon). The samples were then sorted into high and low GFP populations. These populations were then pelleted by centrifugation (4000 rpm for 10 min at 4^0^C) and stored at −80C freezer for downstream analysis. The screens were carried out in biological replicates, under identical conditions, on four different occasions. Sample processing (below) was performed on all samples at the same time to avoid batch effects.

#### Genomic DNA extraction, PCR amplification, and next generation sequencing

Frozen cell pellets were thawed at room temperature. Genomic DNA of the sorted and unsorted fixed cells were then extracted using the protocol described previously^124^. The sgRNA library was amplified by a two-step PCR protocol for NGS^84,125^. One cell consists of approximately 6 pg of DNA and the lowest representation calculated from all samples is 224 x^126^. All gDNA of sorted samples were used to maximize the representations. For the unsorted/bulk samples, 22890 ng of gDNA, which is relevant to 1250 x representation, was used for PCR1. The entire gDNA was amplified using the following primers (NGS-1^st^ PCR Fwd: 5’-TCGTCGGCAGCGTCAGATGTGTATAAGAGACAGggactatcatatgcttaccgt-3’ and NGS-1^st^ PCR Rev: 5’-GTCTCGTGGGCTCGGAGATGTGTATAAGAGACAGgagccaattcccactccttt-3’). Each 100 µL PCR1 reaction contains 50 µL of 2xQ5 Master Mix (New England Biolabs, M0494L), 0.2 µL of MgCl_2_ (stock concentration at 1M), 0.5 µL of each primer (stock at 100 µM), DNA, and water. PCR1 conditions: an initial 5 minutes at 98°C; followed by 35 seconds at 98°C, 30 seconds at 60°C, 45 seconds at 72°C, for 24 cycles; and a final 10 minute extension at 72°C. PCR products from multiple first PCR reactions were pooled and 200 µL was cleaned up for the second step PCR using Nextera XT Index Kit (Lot#10089169). The barcodes used are combinations of N701-712, and S502-508,517. Each 50 µL PCR2 reaction contains 25 µL of 2xQ5 Master Mix (New England Biolabs, M0494L), 5 µL of each index primer (N7xx or S5xx), 62.5 ng of PCR1 product, and water. PCR2 conditions: an initial 5 minutes at 98°C; followed by 35 seconds at 98°C, 30 seconds at 58°C, 45 seconds at 72°C, for 6 cycles; and a final 10 minute extension at 72°C. PCR1 and PCR2 products were purified using AMPure XP Bead-Based Reagent (Beckman Coulter, A63881) according to manufacturer’s instructions.

The samples were sent to OHRI StemCore Laboratories for next generation sequencing where Qubit HS DNA assay was used to measure concentration and Fragment analyzer HS NGS assay (Agilent) was used to assess library fragment size. Sequencing was performed on a NextSeq500 at 150 Cycles High Output 400 million of single-end reads using a 30% PhiX spike-in to control for sequence clustering and diversity. CRISPRCloud2 site was employed to analyse files received from the DNA core. The Enrichment-based screen option was selected. All FASTQ files were uploaded concurrently and assigned to the corresponding groups. After providing all the necessary information, the web browser would initiate the processes of trimming, mapping, and quantifying the sgRNA reads. The processed data were accessed through the link provided.

### Western blot

Whole-cell lysates were prepared by direct lysis with 1× SDS sample buffer. Samples were boiled for 10 min at 95°C and resolved by SDS–PAGE. Briefly, samples were spun down and run on a 6-18% polyacrylamide gel, transferred to a PVDF membrane, and blocked for 15 min with 5 % non-fat milk prior to overnight primary antibody incubation.

### Statistical analysis

Statistical analysis was performed on three biological repeats. Error bars represent the standard deviation in normalized fold changes in observed induction or repression. Statistical significance was determined using two-way ANOVA for three datasets. All statistical parameters for assays in this study are shown in the corresponding figure legends.

### Immunofluorescence

Cells were plated on coverslips 48 hours prior to treatments. After treatments, cells were fixed by 4% PFA in PBS for 15 min at room temperature, followed by permeabilization with 50 μg/mL digitonin (VWR) in PBS for 10 min at room temperature. Cells were blocked in blocking buffer (1% BSA and 2% serum in PBS) for 30 min, then incubated with primary antibodies in the same buffer for 1 h at room temperature. Slides were then washed 2× in PBS and 1× in blocking buffer before incubation with secondary antibodies 1 h at room temperature. Samples were washed 3× in PBS, stained with DAPI (Sigma-Aldrich), and mounted. Images were captured with inverted epifluorescent Zeiss AxioObserver.Z1.

### Quantification of immunofluorescence

A protocol built in the ImageJ software was used to analyse epifluorescent microscopy images to avoid bias. Briefly, channels were first split to examine either number of cells or puncta of interest. The images were changed to 8-bit and set as binary default. Thresholds were then adjusted to identity nuclei or puncta. Finally, particles were analysed and a table summarizing puncta/nuclei quantity and size were provided. The same protocol was applied to each field of view and across samples. Quantification was performed on representative experiments with an average of 9 fields of view per replicate.

## Acknowledgements

The authors acknowledge the support from CIHR grants #153034 (R.C.R) and PJT-169097 (M.W.C.R.) as well as Natural Sciences and Engineering Research Council of Canada #2023-05587 (R.C.R) and RGPIN-2019-04133 and DGECR-2019-00369 (M.W.C.R.) and 201911CGV-434032-74238 (TL). The authors also thank the following Core facilities from the University of Ottawa and the Ottawa Hospital Research Institute (OHRI) for use of their facility, equipment, and expertise: the Cell Biology and Imaging Acquisition Core (RRID:SCR_021845), the Flow Cytometry and Virometry Core (RRID:SCR_023306), the Genome Engineering and Molecular Biology Core (RRID:SCR_022954) and the OHRI StemCore Laboratories (RRID:SCR_012601). We would also like to acknowledge Peter Kim (University of Toronto) for experimental advice on the assessment of pexophagy.

---

**Figure S1: Analysis of autophagy flux in ATG5 KO cells.** The reporter cells were transduced with viruses carrying sgRNA targeting ATG5. These cells were then treated with amino acid-free media for 3 hours, tunicamycin (10 µg/mL) for 6 hours, or clofibrate (1 mM) for 6 hours. Next, they were examined using FACS. Histogram overlays compare either GFP or DsRed signals of the treated ATG5 KO reporter cells with those signals of the untreated ATG5 KO ones.

**Figure S2: Validation of candidates associated with ER stress-induced autophagy.** (A), (B) Wild-type HEK293A cells were infected with both lentiviruses harboring both Cas9 and sgRNA targeting indicated positive (A) or negative (B) regulators. Polyclonal KO cells were then incubated with tunicamycin (10 µg/mL) for 6 hours. Autophagy flux was examined through blots of p62. Experiments were repeated 6 times. p values denote statistical significance of treated KO cells compared to treated control cells and were determined by two-way ANOVA. *p□≤□0.1; **p□≤□0.01; ***p□≤□0.001; ****p□≤□0.0001; ns, not significant.

**Figure S3: Validation of candidates associated with peroxisomal stress-induced autophagy.** (A), (B) Wild-type HEK293A cells were infected with both lentiviruses harboring both Cas9 and sgRNA targeting indicated positive (A) or negative (B) regulators. Polyclonal KO cells were then exposed to clofibrate (1 mM) for 6 hours. Autophagy flux was examined through blots of p62. Experiments were repeated 6 times. p values denote statistical significance of treated KO cells compared to treated control cells and were determined by two-way ANOVA. *p□≤□0.1; **p□≤□0.01; ***p□≤□0.001; ****p□≤□0.0001; ns, not significant.

**Figure S4: Depletion efficiency of KO cells.** Whole-cell lysates of polyclonal KO cells were immunoblotted for the levels of depleted proteins using the antibodies indicated.

