## Supplementary figures and images for "Identification of stress specific autophagy regulators from tandem CRISPR screens"

### Supplemental Figure 1

Screen conditions in ATG5 KO cells

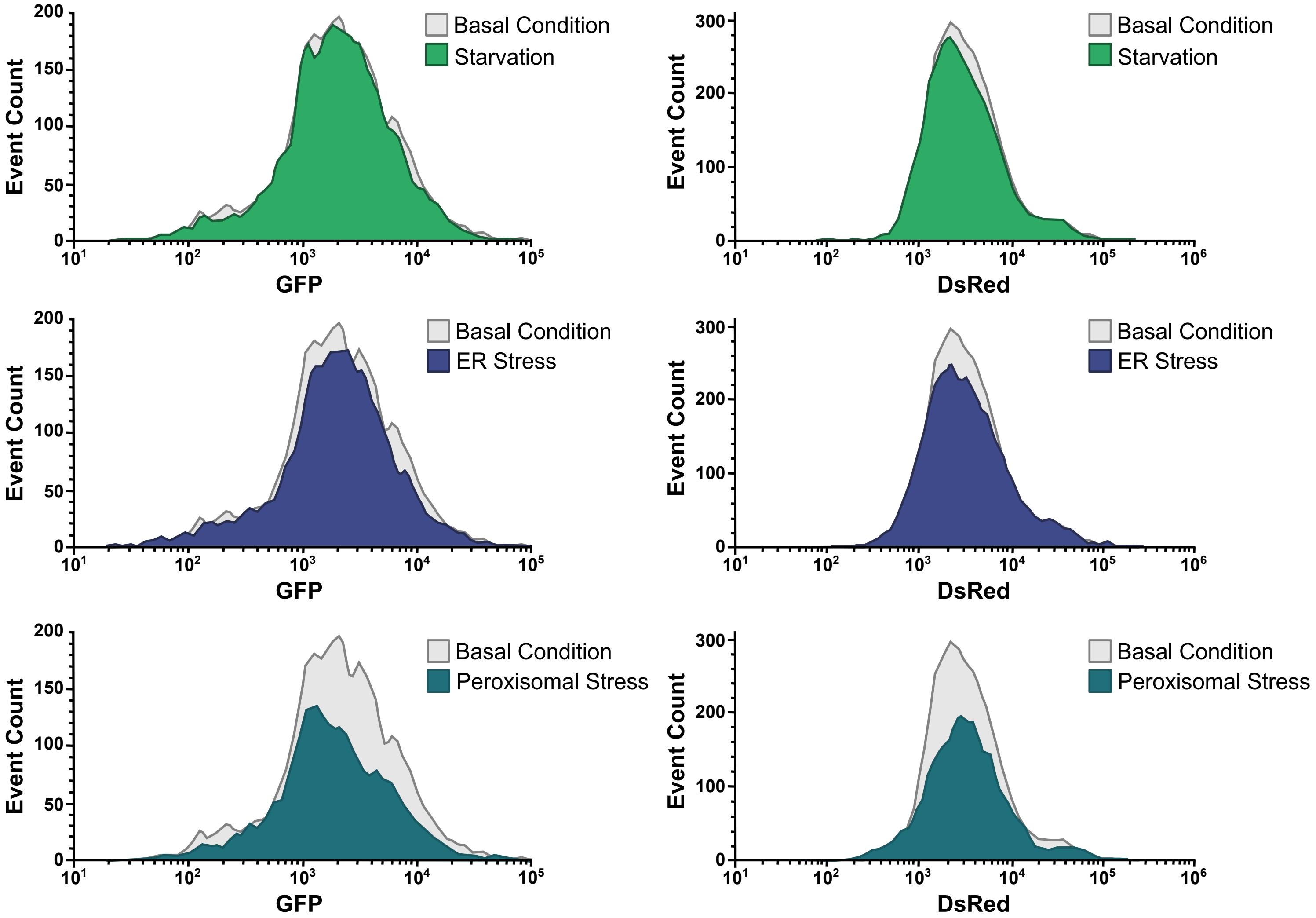


Figure S1

### Supplemental Figure 2

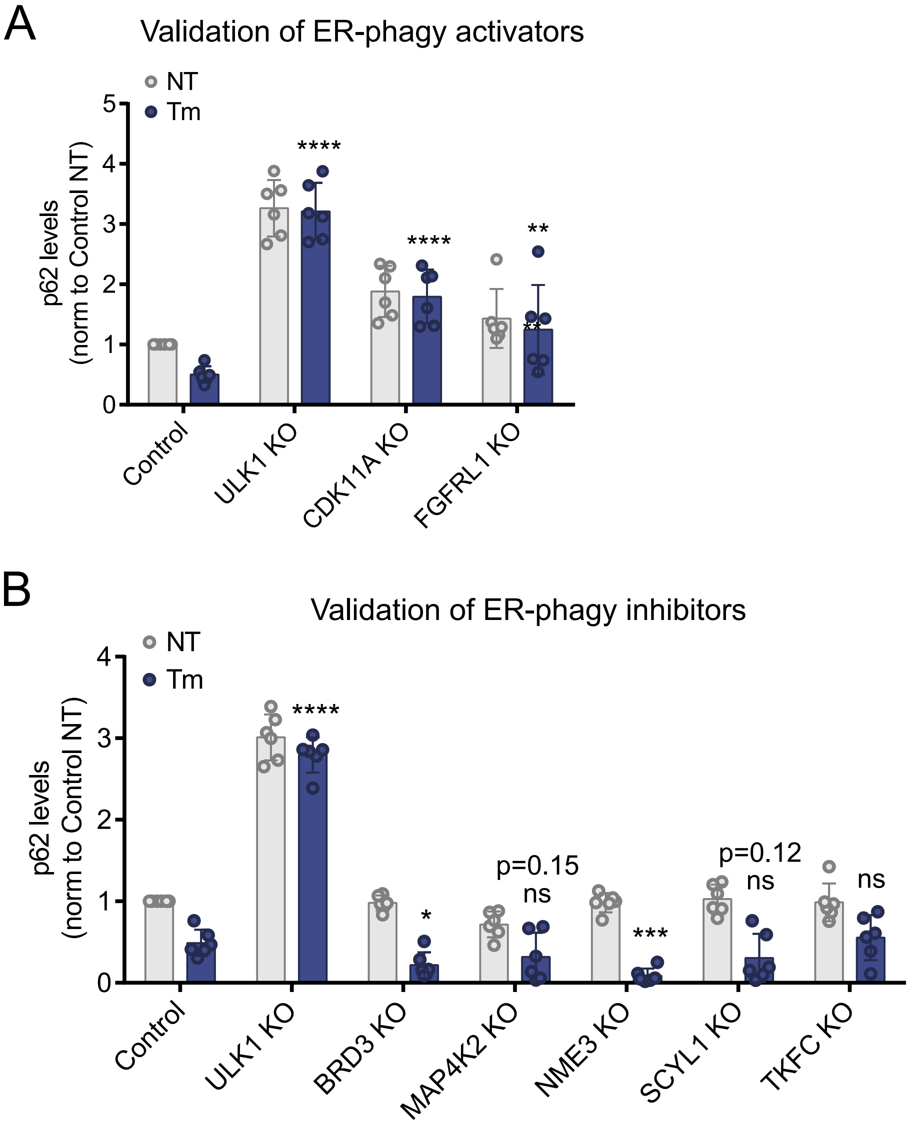

### Supplemental Figure 3

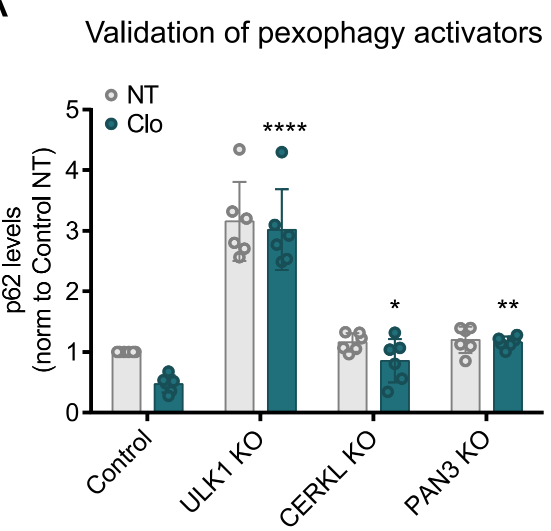

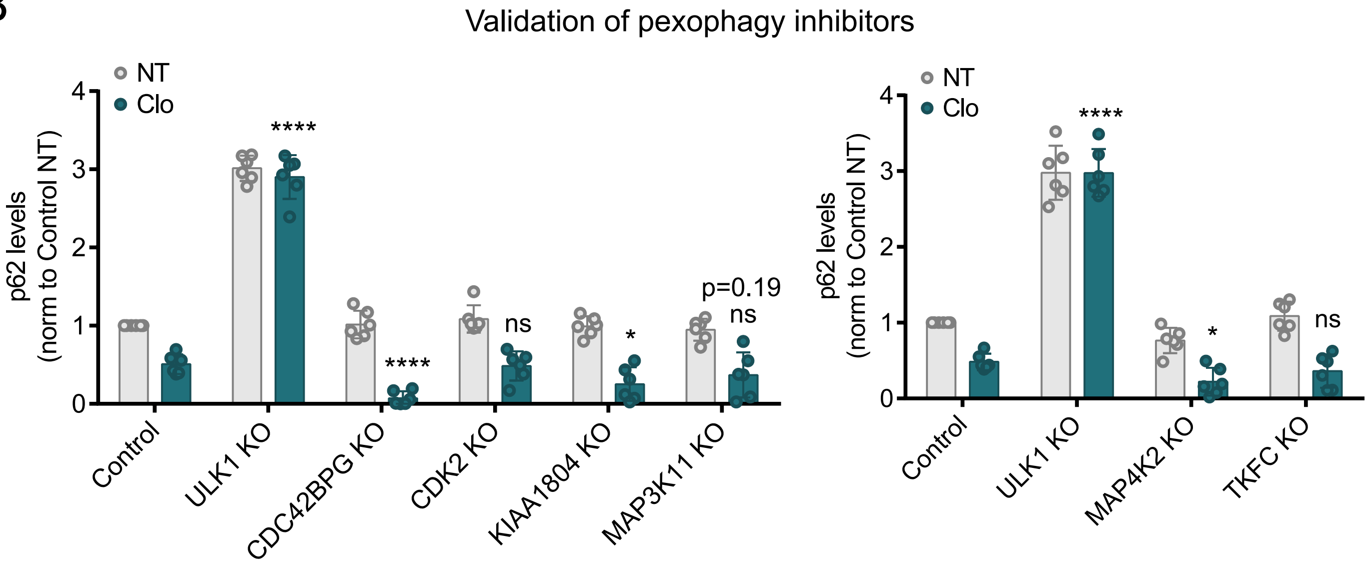

### Supplemental Figure 4

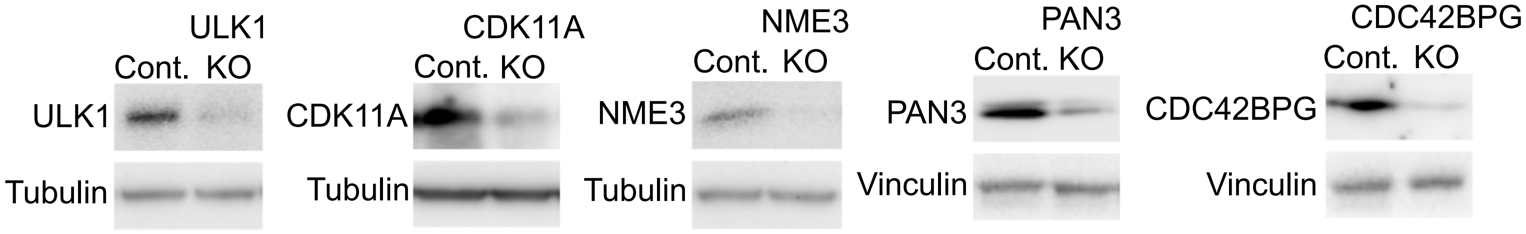
